# Modeling communication and switching nonlinear dynamics in multi-region neural activity

**DOI:** 10.1101/2022.09.13.507841

**Authors:** Orren Karniol-Tambour, David M. Zoltowski, E. Mika Diamanti, Lucas Pinto, David W. Tank, Carlos D. Brody, Jonathan W. Pillow

## Abstract

Understanding how multiple brain regions interact to produce behavior is a major challenge in systems neuroscience, with many regions causally implicated in common tasks such as sensory processing and decision making. However, a precise description of interactions between regions remains an open problem. Moreover, neural dynamics are nonlinear, non-stationary, and can vary dramatically across sessions, days, and animals. Here, we propose multi-region, switching dynamical systems (MR-SDS), a probabilistic model of multiple latent interacting systems that evolve with switching nonlinear dynamics and communication between regions. MR-SDS includes directed interactions between brain regions, allowing for estimation of state-dependent communication signals, and accounts for sensory inputs effects, history effects, and heterogeneity across days and animals. We show that our model accurately recovers latent trajectories, vector fields underlying switching nonlinear dynamics, and cross-region communication profiles in two simulations. We then apply our method to two large-scale, multi-region neural datasets involving mouse decision making. The first includes hundreds of neurons per region, recorded simultaneously at single-cell-resolution across 3 distant cortical regions. The second is a mesoscale widefield dataset of 8 adjacent cortical regions imaged across both hemispheres. On these multi-region datasets, our model outperforms existing piece-wise linear multi-region models and reveals multiple distinct dynamical states and a rich set of cross-region communication profiles.

## 1 Introduction

Advances in neural recording techniques and large-scale electrophysiology (Sofroniew et al., 2016, Song et al., 2017, Liu et al., 2021) have transformed systems neuroscience over the past decade, providing simultaneous measurements of neural activity across regions at high temporal and spatial resolutions. Experiments using these technologies have shown that neural computation is highly distributed (Steinmetz et al., 2019, Cowley et al., 2020, Musall et al., 2018, Allen et al., 2019, Makino et al., 2017, Allen et al., 2017, Gilad et al., 2018). The need for complementary multiregion methods has led to a flurry of activity in this area (Semedo et al., 2014, Kohn et al., 2020, Semedo et al., 2019, Glaser et al., 2020, Perich and Rajan, 2020, Keeley et al., 2020a,b). In this work, we build upon previous contributions in multiregion models, and switching nonlinear time-series models to develop a probabilistic model that accounts for multiregion activity through latent, nonlinear dynamical systems that evolve and communicate in a direction fashion at each timestep. The approach models high dimensional multi-region observations as emissions from coupled, low dimensional dynamical systems with explicit local dynamics and communication. We additionally introduce a measure of the volume of communications between brain regions, allowing us to quantify the directional ‘messages’ communicated between regions across time.

## 2 Background

### 2.1 Nonlinear neural dynamics

A wide variety of models of sequential neural data with nonlinear dynamics have been introduced in recent years (Linderman et al., 2017, Pandarinath et al., 2018, Duncker et al., 2019, Kim et al., 2021). One approach is to model nonlinear dynamics as a combination of linear regimes Linderman et al. (2017), Nassar et al. (2019), Zoltowski et al. (2020). While these switching linear models can generally represent nonlinear dynamics, they require multiple discrete states to instantiate a single nonlinear dynamics vector field, which reduces interpretability. Similarly, the expressivity of each linear regime depends on the number of latent dimensions used, which can make visualization challenging. A separate approach is LFADS, which uses a sequential autoencoder with a generative recurrent neural network (RNN) (Pandarinath et al., 2018). While this approach is powerful, the RNN typically must be high-dimensional to learn complex dynamics, which makes visualization of low-dimensional dynamics vector fields challenging. Additionally, the model only instantiates a single set of nonlinear dynamics, whereas our aim is to learn multiple nonlinear dynamical regimes. Recently, Kim et al. (2021) introduced a powerful model, PLNDE, that uses a neural ODE to model nonlinear, low-dimensional dynamics in continuous time. We view our proposed method as a complementary discrete-time approach that enables learning multiple nonlinear dynamics vector fields.

### 2.2 Switching nonlinear dynamics

Two recent papers introduced switching nonlinear dynamics models. First, Dong et al. (2020) proposed a model with switching nonlinear dynamics that uses an amortized inference network to infer continuous latents, and developed a collapsed variational inference approach for the discrete latent variables. Next, Ansari et al. (2021) proposed to explicitly model the duration of discrete latent occupancy through switch time step counts. They showed the semi-Markov formulation allowed for improved inference of long-duration states. The approach taken in this paper builds on the work of both of these papers, generalizing SNLDS and RED-SDS Dong et al. (2020), Ansari et al. (2021) to multiregion neural data with inputs.

### 2.3 Multi-region modeling and communication

Various approaches to modeling multi-region neural data have been proposed (Semedo et al., 2014, Kohn et al., 2020, Semedo et al., 2019, Glaser et al., 2020, Perich and Rajan, 2020, Keeley et al., 2020a,b). Our model directly relates to mp-rSLDS, the multiregion recurrent switching linear dynamical systems model introduced in Glaser et al. (2020). In this setup, the observation process is constrained such that continuous latent states map uniquely to observations originating in a particular brain region. MR-SDS can be seen as a generalization of mp-rSLDS to a model with nonlinear dynamics, communication, and emissions.

### 2.4 Decision making dynamics

Decision making processes have been modeled extensively as variants of a bounded drift-diffusion process, in which evidence-driven drift is combined with a noise diffusion process to produce accumulation, or a ‘race’ between competing evidence sources towards a decision boundary (Gold and Shadlen, 2007, Ratcliff and McKoon, 2008, Brunton et al., 2013, Pinto et al., 2018, Koay et al., 2019). A common paradigm for studying decision making is sensory evidence accumulation, in which an animal must accumulate competing sensory cues towards a decision (Brunton et al., 2013, Latimer et al., 2017, Mazurek et al., 2003). Two emerging experimental results add complexity to understanding the neural dynamics underlying decision making. The first is that the neural basis of decision making appears to be highly distributed. Recent work has shown evidence and decision related information is present in many brain regions (Pinto et al., 2020, 2019), and precise inactivation studies have provided causal evidence for the necessity of multiple regions in the accumulation process (Pinto et al., 2020). However, the precise role of different regions in the accumulation and decision process remains an open problem, underscoring the need for novel multiregion analysis methods that can help address questions about local computation, cross-region communication, and stimulus effects. The second challenge is that decision-making behavior appears to exhibit persistent states in which different decision strategies are employed (Ashwood et al., 2020, Stone et al., 2020). This indicates the neural dynamics underlying decision making may not be consistent within and across trials, motivating our use of switching nonlinear dynamics. Finally, recent work has shown cue-locked responses in early visual areas and intermediate brain regions are modulated by recent sensory stimuli and evidence history (Koay et al., 2019). These findings motivate our use of nonlinear interactions between brain regions, history dependence, and stimuli effects.

## 3 Multiregion, switching dynamical model

### 3.1 Switching nonlinear dynamics and communication with multiple latent regions

Here we describe the multi-region, switching dynamical systems (MR-SDS) model in detail. The model is designed to capture the following key features. First, our goal is to learn low-dimensional, nonlinear dynamics and corresponding latent trajectories underlying neural data. We expect that neural dynamics are nonstationary, and that different nonlinear vector fields may be better descriptions of neural dynamics at different timepoints. We therefore incorporate a discrete state that enables switching between different sets of nonlinear dynamics. Secondly, we are primarily interested in modeling multiregion neural data, and thus our model is designed to separate the activity of each brain region into different continuous latent dimensions, such that the within-region and across-region dynamics are accessible for analysis. Finally, we desire to include a nonlinear mapping between the continuous latent and observed neural data, to reflect the nonlinear observation process of calcium imaging.

### 3.2 Discrete switches between different dynamical regimes

The MR-SDS model has both continuous and discrete latent variables, respectively denoted by x and *z*. The generative process is

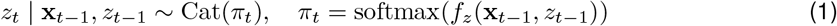

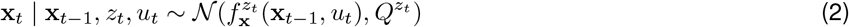

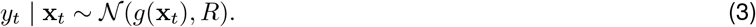

Above, the discrete state *z*_*t*_ switches probabilistically at each timestep as a function of the continuous latent x, as well as its own history (similar to a Hidden Markov Model or HMM). It is also possible for the inputs *u*_*t*_ to effect transitions directly. The discrete state *z*_*t*_ in turn indexes the dynamical regime active at time *t*, with different dynamics 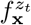 governing the continuous latent variable x_*t*_ for each discrete state. Hence, changes in the discrete state over time cause switches in the nonlinear dynamics. Notably, the model does not explicitly specify a condition or distribution on switching locations in state space or switching times induced by discrete transition dynamics *a priori*; the transitions are instead probabilistic and not known in advance. Thus, the model is free to learn transitions driven by a combination of continuous and discrete states, and possibly external inputs, without constraints.

### 3.3 Multiregion dynamics and emissions structure

Importantly, we constrain the model to have a multi-region structure. We consider a decomposition of the global continuous state into *K* variables private to each region 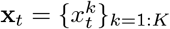, and define the overall global dynamics function 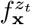 using additive components acting on each region’s latents 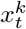. Thus, the dynamics function for the *k*’th region is

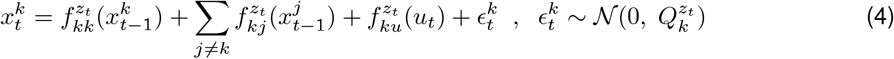

The corresponding emissions *g* for the *k*’th region are defined as

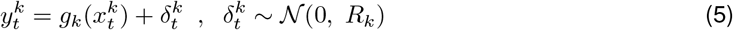

Above, 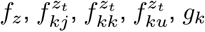 are nonlinear functions parameterized by neural networks. The generative model accounts for each region’s observations *y*_^*k*^_ as nonlinear ‘private’ emissions from a corresponding continuous latent variable *x*_^*k*^_. *z*_*t*_ remains a global switch which indexes which of *M* local dynamics and communication functions are active at each timepoint. Thus, 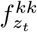 indicates the nonlinear local dynamics indexed by for region *k* active at time *t*, indexed by discrete state *z*_*t*_. Note emissions in the model are constrained such that latents from one region only map to that region’s observations. This model is thus a generalization of multiregion LDS-based models such as mp-rSLDS (Glaser et al., 2020). For spiking data, it’s also possible to use the emissions mapping 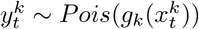.

### 3.4 Estimating cross-region communication and input effects

By explicitly inferring each region’s local dynamics, communication and input processing functions, the model exposes estimates of communication streams between regions. We define “messages” and regional input effects in the model follows:

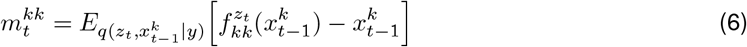

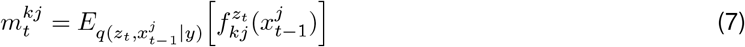

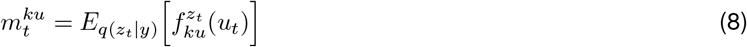

Where above the expectation is taken with respect to the approximate posterior over the latent states, *z* and *x*. Taking the expectation requires explicitly marginalizing over *q*(*z*_*t*_), while we can directly take the mean of the Gaussian given by *q*(*x*). To quantify the message volume, we use the Frobenius norm 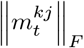. Thus, the model allows us to produce estimates and quantify communication and input effects for further downstream analysis (eg Figure 5).

**Figure 1:**
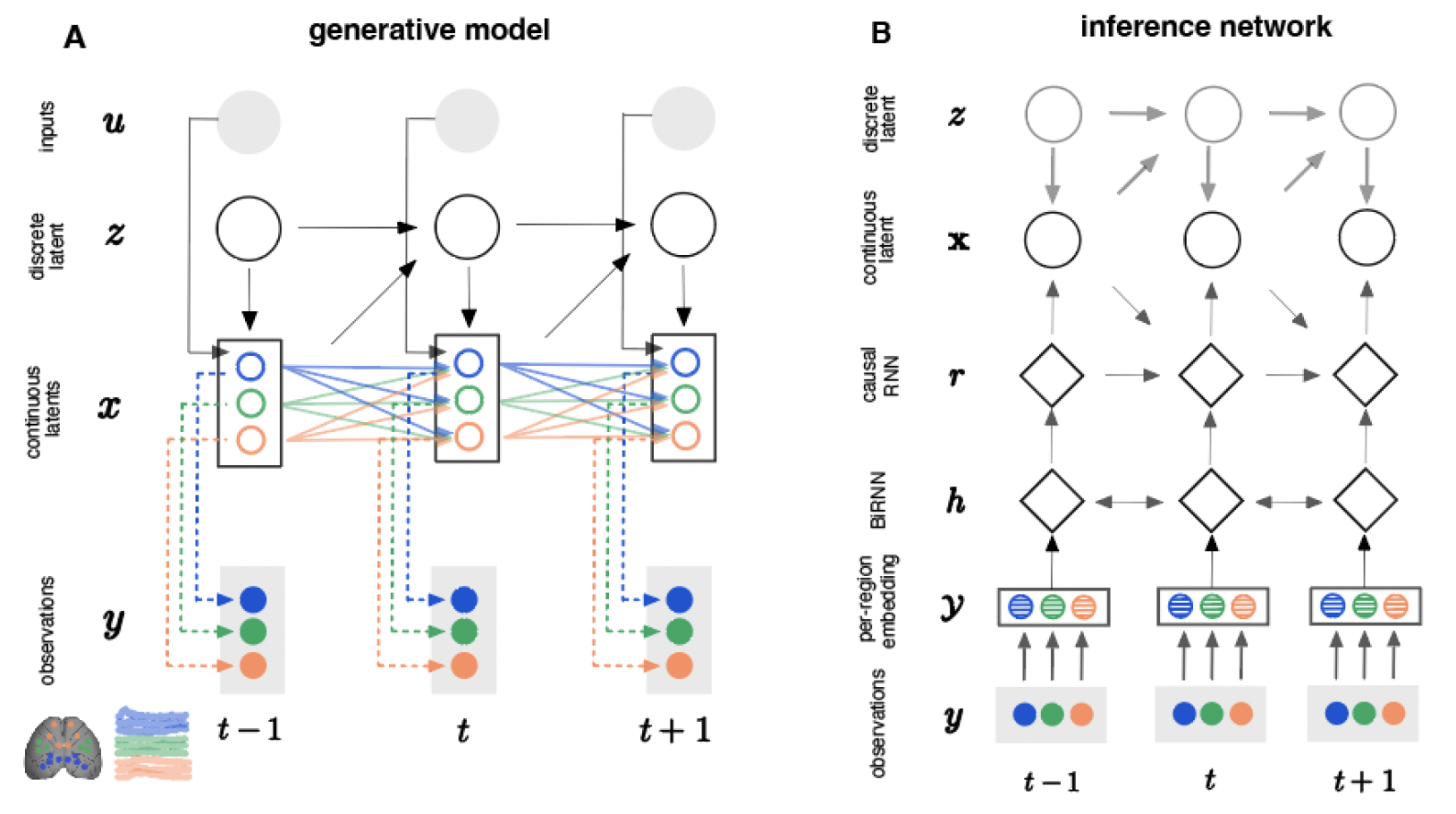
Generative model and inference network.

**Figure 2:**
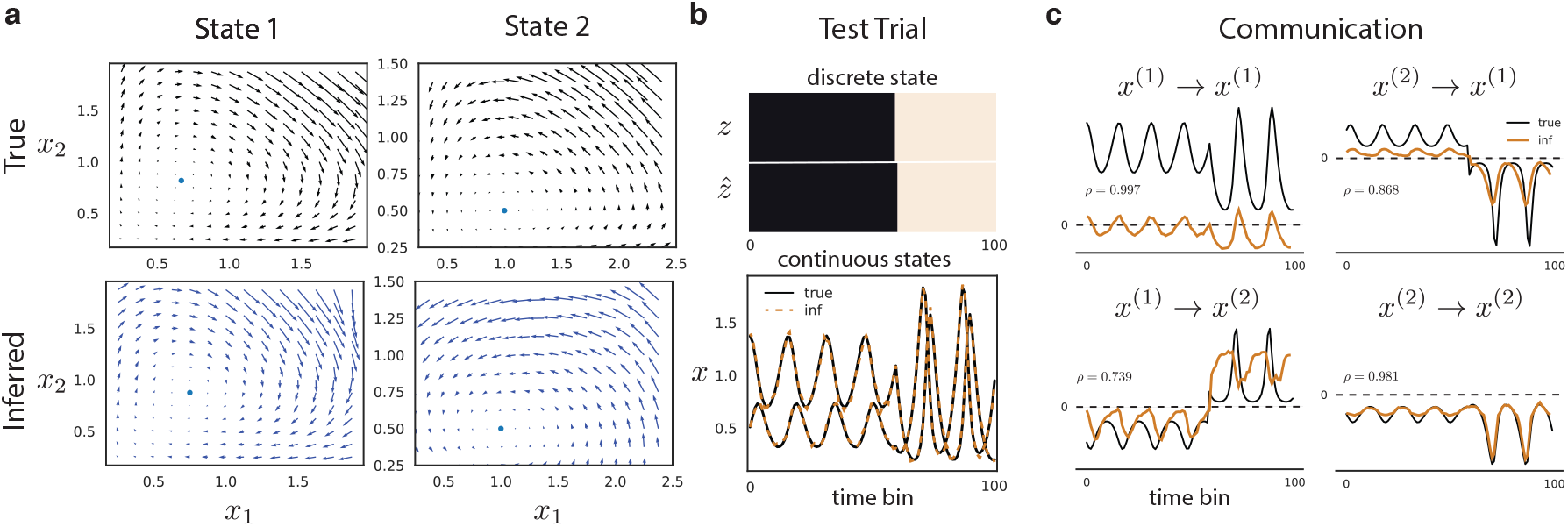
Multi-region Lotka-Volterra simulation. **(a)** True and inferred dynamics vector fields and fixed points across two discrete states. The inferred dynamics were mapped with a linear transformation to account for non-identifiability. **(b)** True and inferred discrete and continuous latent variables. The continuous latent variables were mapped via a linear transformation. **(c)** True and inferred communication profiles within and between regions.

**Figure 3:**
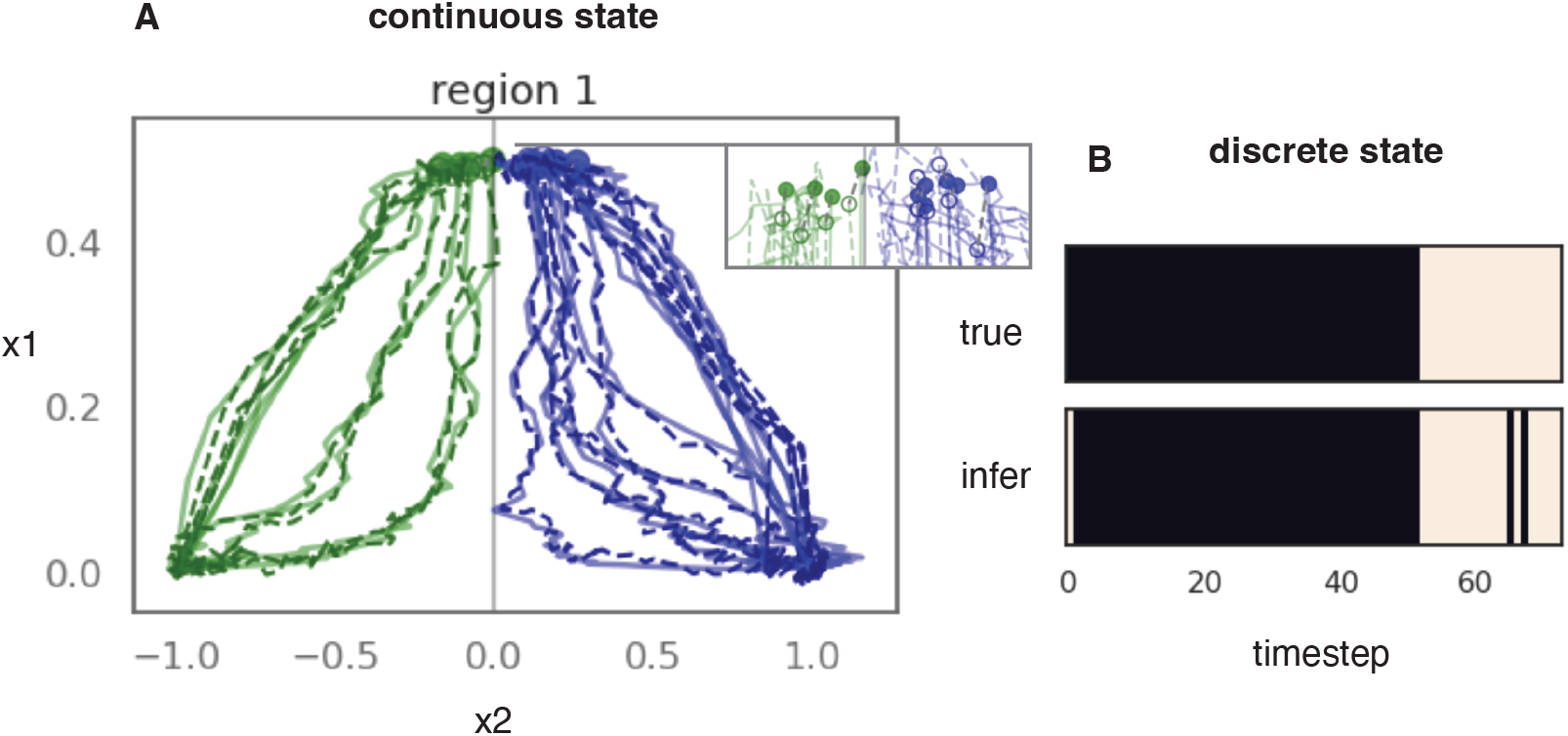
Multi-region, hierarchical evidence accumulation simulation with double well dynamics. Region 1 (accumulator) shown. A. Recovery of continuous latent trajectories for a series of left (green) and right (blue) trials; true trajectories plotted with dashed lines. Inset: recovery of initial condition for each trial, true initial condition denoted with open circle. B. Recovery of discrete states shown for a single representative trial.

**Figure 4:**
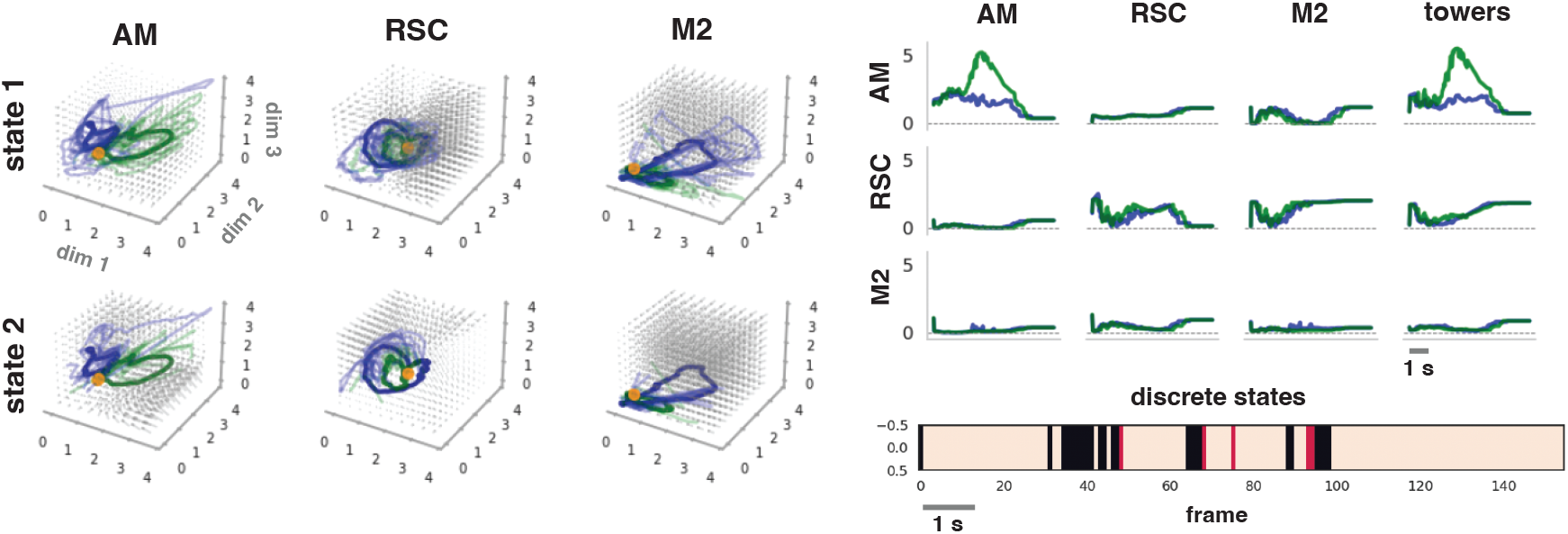
Multiregion mesoscope imaging data. Left: inferred dynamics vector fields and trajectories for left (green) and right (blue) trials on test data. Average trajectories overlaid in bold, starting location in orange. Right: Inferred message norms from sending (columns) to receiving (rows) for left and right trials. Note left (in green) effect in visual area AM.

**Figure 5:**
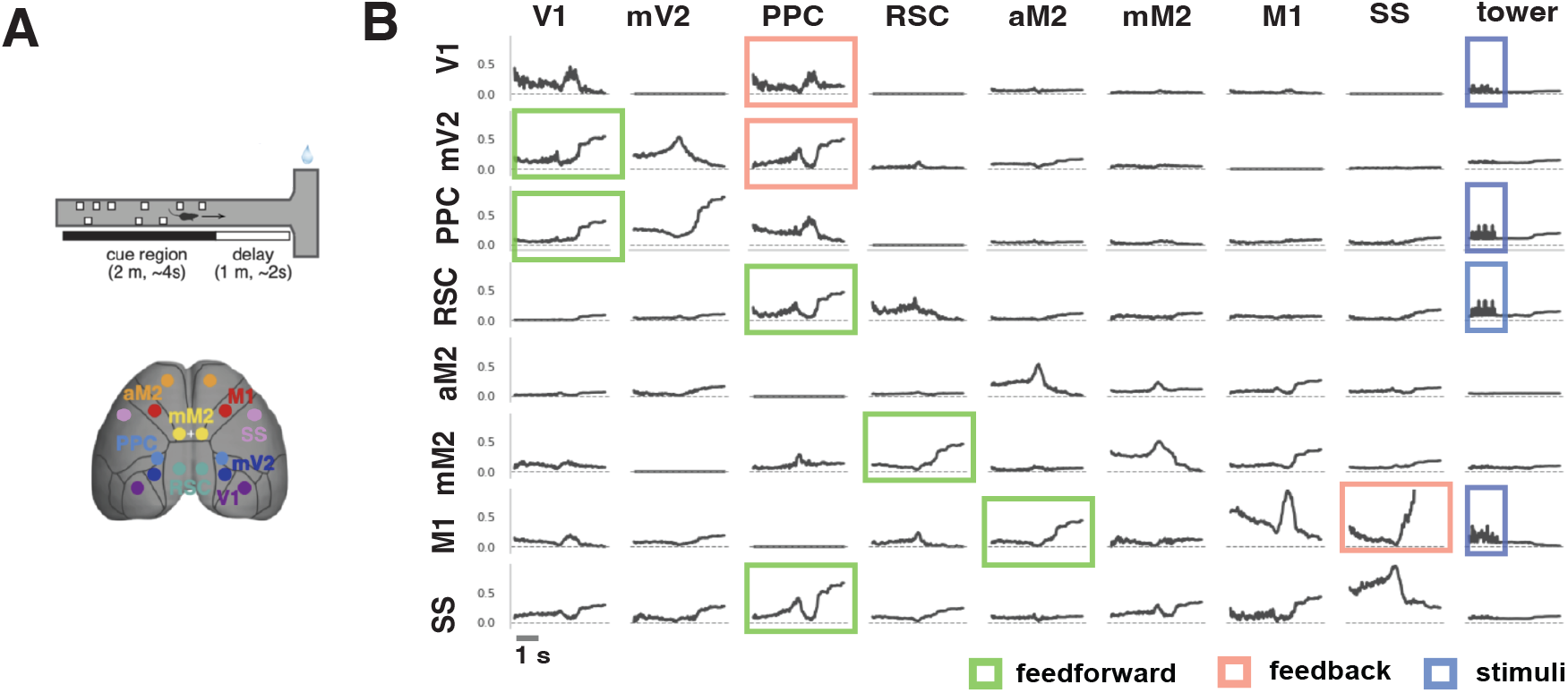
Multiregion mesoscale widefield imaging data. A. Illustration of the towers task; imaging was performed over 8 regions across both hemispheres. B. Macro view of communication streams. Trial averaged inferred message norms going from each sending (columns) to receiving (rows). Highlighted: communication streams with notable feedforward, feedback, and stimuli effects.

### 3.5 Optional dependence of input processing and communication on local state

There is growing evidence that stimuli effects and communication depend on local activity in the target region Koay et al. (2019), Pinto et al. (2019). Conditioning on local activity opens the possibility of capturing local gating or modulation of stimuli and communication. The dynamics for the *k*’th region in this variant take the form:

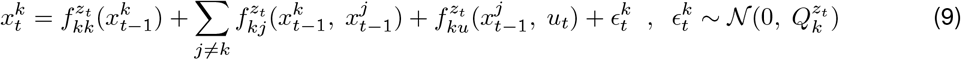

Note the dependence on 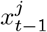 in both incoming messages and input effect terms. The dynamics are still additive, but communication and stimuli effects are conditioned on the local continuous latent state. Thus, we extract locally conditioned messages by comparing to the case in which inputs are zero:

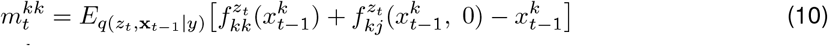

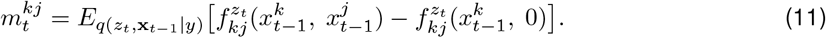

Importantly, the sum of the message components is equal to the effect of the overall transition function for the continuous latents of each region, i.e. 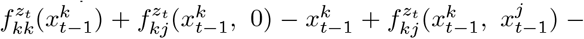 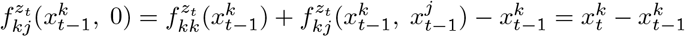.

### 3.6 Optional explicit duration counts on discrete state switching times

Experimentalists may wish to capture strong priors on timing and persistence of discrete states, corresponding to experiment structure. Following Ansari et al. (2021), we condition the discrete state switches on an explicit duration variable *ct*. The augmented model is:

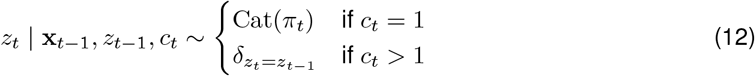

Above, 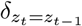 denotes the delta function, taking value 1 when the condition is true. Conditioning on the count persists the current state when the count *c*_*t*_ is above 1; otherwise transitions are sampled from the transition distribution as before. Additional detail is presented in the appendix. Notably, the model without explicit duration is a special case of the explicit duration model, with support on a count of 1 (Ansari et al., 2021). **Figure A4** shows the explicit duration model infers longer discrete switches on data, underscoring the potential importance of modeling switch times explicitly.

### 3.7 Capturing trial history effects with a per-region, conditional initial condition

To account for trial history effects, we condition each region’s initial condition on the previous trial’s observations for that region, choice, and reward, as follows:

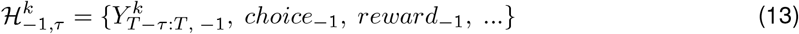

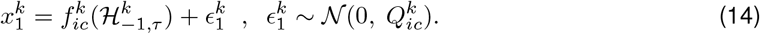

### 3.8 Amortized inference with multiregion structured encoder network

Each region’s observations are embedded by a region specific sub-network. The embeddings for all regions are processed by a shared RNN backbone that predicts the next latent x.

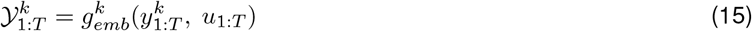

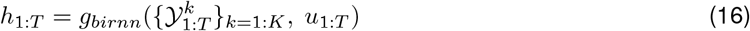

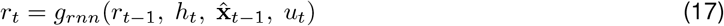

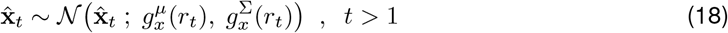

Above 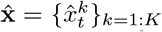 is the sampled estimate of x. 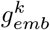 is a feed-foward neural network embedding region *k*’s activity 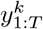 along with observed stimulus inputs *u*1:*T* · *g*_*birnn*_ and *g*_*rnn*_ are a non-causal, bidirectional RNN and a causal RNN,, together forming the shared inference backbone. Finally, 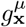 and 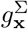 are feed-forward networks mapping *r*_*t*_ to the mean and covariance of the inferred latents at time *t*. Following (Dong et al., 2020, Ansari et al., 2021), we treat samples 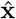 from the approximate posterior q(x) as pseudo-observations for the subgraph 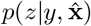, which forms an HMM with *z* as discrete states and 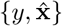 as the emissions. This involves estimating the approximate posterior conditioned on the data, with the exact posterior conditioned on the samples 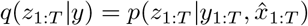. Inference for *z* can be done in closed form using the forward-backward (FB) algorithm (Rabiner, 1989), which we also differentiate. In the explicit duration case, introduction of a learned distribution over duration counts makes the discrete transition model *z*_*t*_ | _*zt−*1_, *x*_*t−*1_, *c*_*t*_ a semi-Markov model. The discrete states can then be estimated using the FB algorithm for HSMMs (Hidden Semi-Markov Models) (Ansari et al., 2021, Chiappa et al., 2014).

## 4 Experiments

All experiments and analysis were run on a 28 CPU, 8 GPU (GeForce RTX 2080 Ti) server.

### 4.1 Multi-region switching Lotka-Volterra simulation

We first demonstrate the multi-region SDS modeling approach in a simulated example that switches between two different sets of nonlinear Lotka-Volterra dynamics with multi-region observations. Conceptually, our intention is to emphasize how learning switching nonlinear dynamics vector fields can uncover interpretable changes in dynamics over time. Additionally we demonstrate that the per-region inference methods are effective. The true simulated dynamics switch between two sets of Lotka-Volterra dynamics

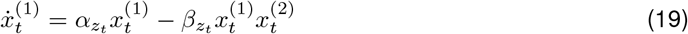

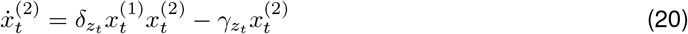

where each dynamics parameter depends on the current discrete state *z*_*t*_ (Figure 2). Lotka-Volterra dynamics are a two-dimensional model of interacting predator-prey populations Lotka (1925); we model each latent dimension as a separate region, such that half of the observations are a function of *x*_1_ and half are a function of *x*_2_. This is an example with oscillatory coupling, one potentially important mode of neural communication (Kohn et al., 2020, Kastner et al., 2020). The model was simulated in discrete time using the Euler approximation. The simulated trials were 100 time steps long; for each trial, a discrete state switching time was sampled uniformly between 40 and 60. Each discrete state corresponded to a different set of dynamics given by parameters 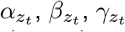 and 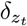, with index *z*_*t*_ indicating their value changes based on the discrete state. Figure 2a shows dynamics for the two states on both dimensions of the joint state space. The model had linear observations, a multi-region embedding network, and was fit with *M* = 2 discrete states, with a global set of nonlinear dynamics in each state

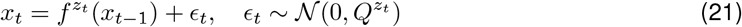

Where above, 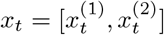. Learned dynamics under the fitted model matched the underlying true dynamics, as shown by similarity in the dynamics vector fields and fixed points (Figure 2a). Additionally, estimates of discrete and continuous latent states on simulated test trials closely matched the known ground truth (Figure 2b). Next, we computed true and inferred communication profiles. Following equations (10, 11), we computed the within region dynamics as 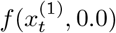 and across region dynamics as 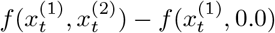. Importantly, these two terms sum to the overall dynamics function *f*. The profiles are marginalized over the discrete states based on discrete state posteriors. The true and inferred communication profiles over time were highly correlated (Figure 2c). We did not compute a transformation between true and inferred profiles. Importantly, inferred across-region communication profiles correctly change in sign when the discrete state switches.

### 4.2 Double-well hierarchical accumulation dynamics

To further motivate the use of our model for neural data, we simulated a multiregion system performing a decision-making task. A recent finding is that animals performing accumulation in sequential trials appear to display robust and structured sequences of behavior, including long periods of lapses, bias and strategy switches (Ashwood et al., 2020, Stone et al., 2020, Koay et al., 2019). Thus, our aim was to show how intrinsic dynamics of a multiregion, switching nonlinear model can account for accumulation dynamics across consecutive trials in a way that produces history effects and bias; and to show our model recovers the key aspects of such a system. The system was composed of two regions, an ‘accumulator’ (region 1, or R1) and ‘controller’ (region 2, or R2). R1 is presented with competing stimuli and accumulates. R2 observes dimension *x*_1_ of R1, providing information about the progression of the decision process, but not the ‘winning’ side. When R2 latent activity reaches a transition border, it triggers a dynamics state switch in both regions. R2 then drives feedback to the accumulator, pushing R1 back approximately to its starting condition. This final location serves as the initial condition for the next trial. R2’s internal variability causes variability in initial conditions, leading to biased trial sequences, potentially similar to those seen in animals (Ashwood et al., 2020, Stone et al., 2020). We provide simulation details including system equations and figures in Appendix A. In Figure 3, we show that the model accurately recovers example trajectories simulated from this model.

### 4.3 Evaluating performance on neural datasets

To evaluate the performance on neural data, we compared held out model predictions for a family of related multi-region latent variable models (Table 1). These include PCA, LDS, SLDS, and rSLDS. Importantly, these models can be seen as special cases of MR-SDS, instantiated with less expressive components: PCA has no dynamics and linear emissions; rSLDS has linear transitions, dynamics and emissions. We used the ‘co-smoothing’ metric (Macke et al., 2011, Pei et al., 2021), which evaluates reconstruction loss on 25% held-out neurons in held-out trials. Cosmoothing requires infering latents at test time based on a subset of neurons, and predicting activity of a held out set from those latents. The latent representation thus has to be robust to input removal, a more challenging test than simple reconstruction. MR-SDS model better performance than comparable models on both datasets. We additionally compared to rSDLS models with higher latent dimensionality, observing that mr-rSLDS models required 10 latent dimensions per region (vs 3) to achieve comparable performance on the mesoscope dataset. Finally, we found MR-SDS was comparable with rSLDS models fit to the entire dataset as a single region, providing a baseline for performance without multiregion description.

**Table 1:**
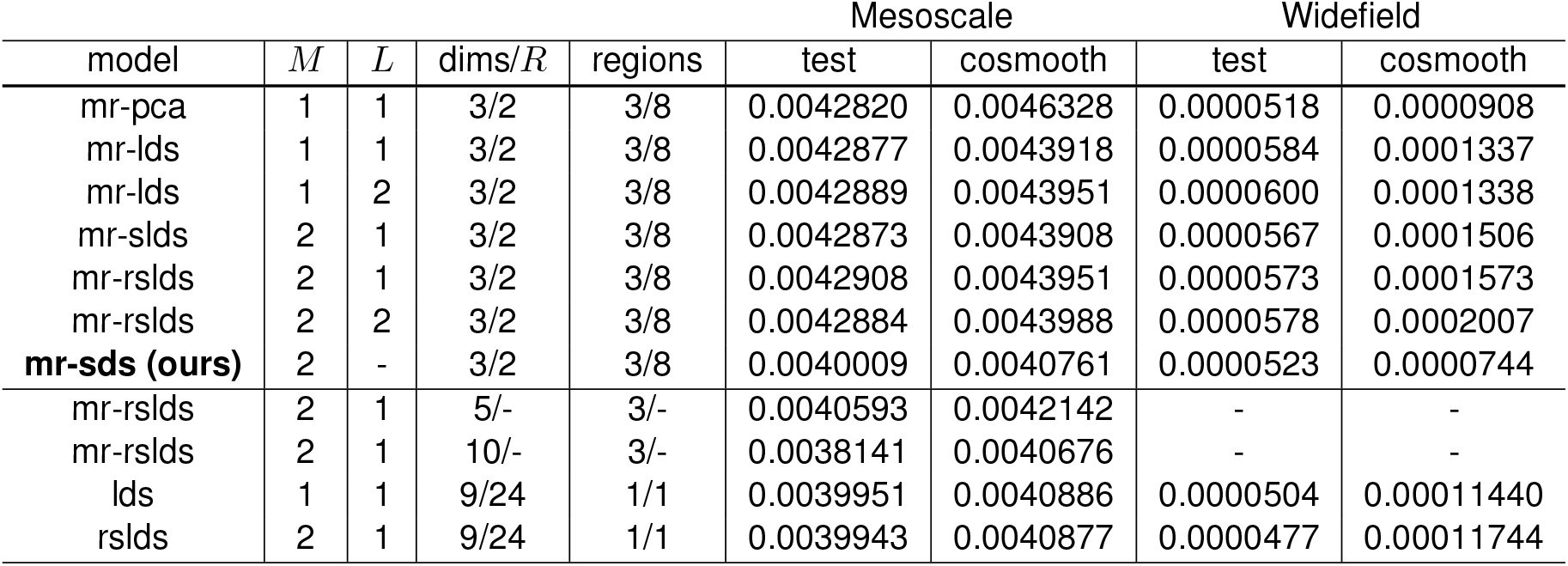
Held-out test error across models on the mesoscope and widefield datasets. *M* : number of discrete states. *L*: number of time-lags.

### 4.4 Application: 3 region, single-cell resolution mesoscope data

We applied our method to calcium imaging data recorded in mice performing a sensory evidence accumulation task. In the task, a headfixed mouse runs on a linear track in virtual reality while columns (‘towers’) are presented on presented on both sides of the track (Pinto et al., 2019). The mouse must decide on which side more towers were presented and turn to correctly to get a reward following a short memory delay section in the track. We analyzed a single day of mesoscope calcium imaging data of 3 distant brain regions in a single hemisphere, consisting of 178 correct trials. The regions were: AM (a visual area) retrosplenial cortex (RSC), and M2 (a higher level motor / planning area). Preliminary results showed distinct trial averaged communication profiles for left and right trials, as well as strong stimuli effect for visual area AM. Further analysis may focus on the state occupancy across trials, and further analyze the interaction between communications and the inferred dynamics vector fields. Additionally, decoding analyses from the inferred messages may be used to quantify the information present in these messages.

### 4.5 Application: 8 region, mesoscale widefield data across hemispheres

We also applied our method to wide-field calcium imaging data from the same experimental task. 16 regions (8 bilaterally) were imaged across the cortical surface (Figure 3). We analyzed a single day of recordings consisting of 63 correct trials. To analyze communication between regions, we examined estimated messages under the inferred model parameters and approximate posterior over the continuous latents. To quantify the messages, we computed their Frobenius norm at each timepoint. Figure 5 shows a ‘macro’ picture of communication streams between all regions. Highlighted are streams with notable feedforward, feedback and stimuli components. We draw attention to a few important features. First, profiles showed strong feedforward drive from V1 to mV2 and PPC. Second, PPC showed strong feedback drive back to V1 and mV2, as well as feedforward drive to upstream regions RSC and SS. These results are potentially consistent with a hypothesized central role of PPC in evidence accumulation, suggesting PPC may act as a hub between early visual areas and upstream regions (Pinto et al., 2019). Further analysis may seek to probe the timing of single trial communications, as well as their relationship to evidence levels in the trial.

## 5 Discussion

We proposed a switching, nonlinear dynamics approach for modeling multi-region neural data. An important question is what benefits switching nonlinear dynamics provides over non-switching nonlinear dynamics. Our answer is that the switching dynamics enables modeling the dynamics in low-dimensional space using more persistent discrete states. This is in contrast with rSLDS models, which must use multiple discrete states to implement a single set of nonlinear dynamics, and alternative approaches such as RNNs that use higher-dimensional spaces to capture the full dynamics range. We validated our proposed model and inference approach in multiple simulations and demonstrated it on two multiregion datasets. There remain important aspects of future work. In particular, our method does not allow for unobserved regions or neural activity to influence dynamics of the observed regions. The learned interactions in our model are also not causal. Additionally, future work should explore alternative types and parameterizations of multiregion communication, and demonstrate whether our approach succeeds or fails in each case.

## Acknowledgement

We thank Brian DePasquale for helpful discussions and his formulation of a double well accumulation energy function that we modified for the double well simulation.

## A Appendix

### A.1 Cosmoothing test time evaluation

A key question in evaluating unsupervised models of neural data is regarding the appropriate metric to use to assess model fits. A limitation of looking only at reconstruction error on held out data is that the model may simply learn to compress and decompress the observations, effectively learning an identity mapping. This can be seen in the relative strength of the PCA benchmark (more on this in MR-PCA section below) on test reconstruction alone. An alternative suggested by (Macke et al., 2011) and adopted in the recent Neural Latents Benchmark (Pei et al., 2021), is “co-smoothing”, in which a model is evaluated on reconstruction loss of held-out neurons on held-out trials (see Figure A1). This tests the ability of the model to learn a latent representation of the data that is robust to removal of inputs, and is thus a more challenging test of a model’s performance. A key result in the benchmarks is that models that perform reasonably well on test reconstruction may perform poorly on cosmoothing. The Neural Latents Benchmark uses a 25% neuron drop out rate, and we apply this rate, with one modification - we evaluate all models against a 25% drop out rate per region. This ensures that results are not dominated by regions with more neurons.

### A.2 Cosmoothing multiregion dropout training

To improve model generalization and to allow it to accurately infer latent variables given only a subset of neural responses, we trained the model on real neural data by dropping out a subset of inputs to the inference network in each batch. In particular, we dropped out inputs from individual neurons over time. Notably, it was important that we drop out equivalent fractions of neurons from each brain region.

To make MR-SDS and the latent represention it learns robust to missing inputs inherent in the co-smoothing test, we modify the training procedure as presented in algorithm 1.

There, the monte carlo samples used to evaluate the expectation are taken with respect to the approximate posterior evaluated on the dropped inputs. We dropped out 25% of neurons per region. We found that dropout of 50-80% of trials works well, depending on the dataset. Empirically, datasets with higher correlation between observations, such as the widefield dataset used in the experiments, require a lower trial dropout rate. We note that this is similar to the ‘coordinated dropout’, or ‘speckled’ holdout strategy used by (Keshtkaran and Pandarinath, 2019), but extended here to multiregion data, and we drop out all of the timepoints for random neurons on each trial.

#### Algorithm Cosmoothing multiregion drop out training

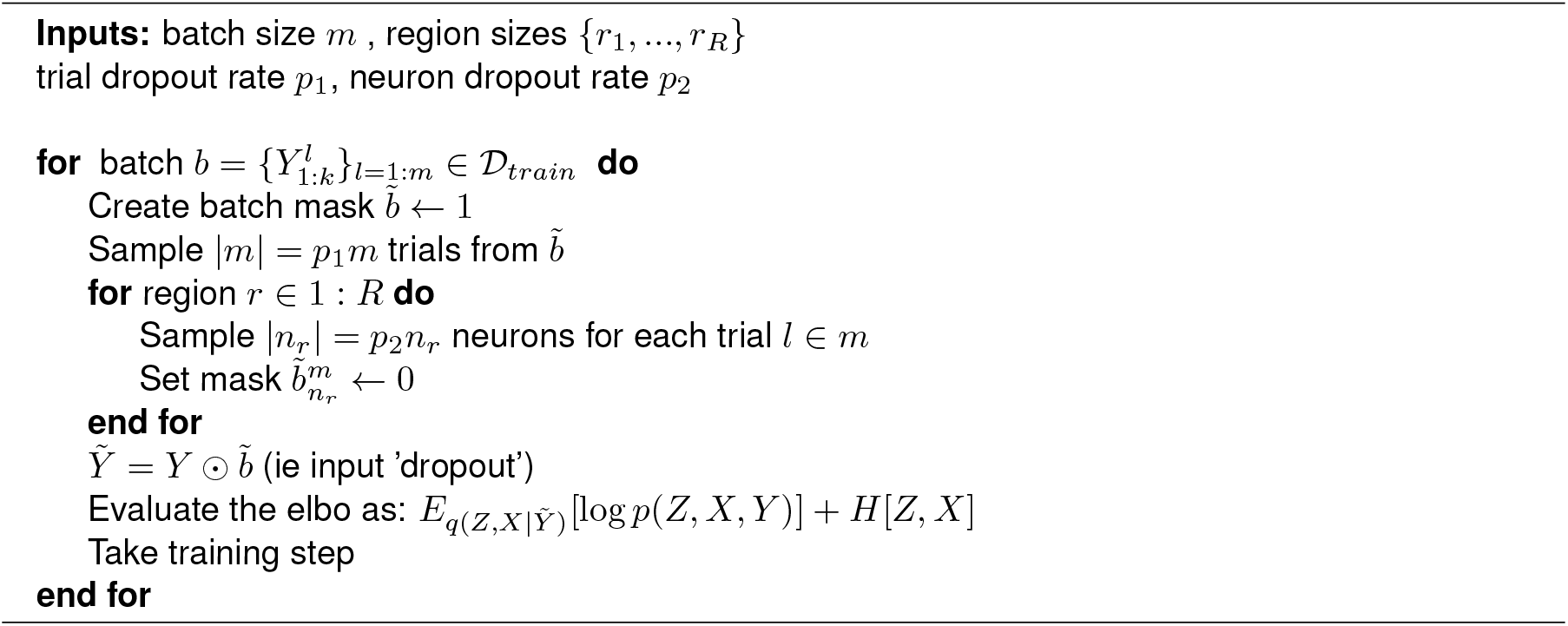

We used a dropout rate *p*_1_ = {0., 0.5} for the mesoscope and widefield datasets respectively in the experiments presented in the paper.

**Figure 6:**
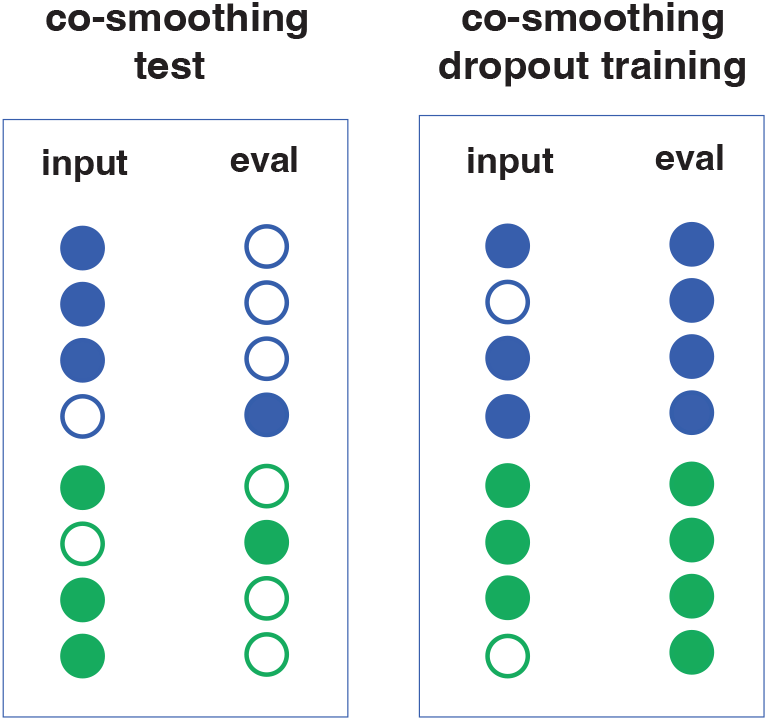
Multiregion cosmoothing and multiregion cosmoothing dropout training. **Left:** When cosmoothing, we drop a percent of neurons from each region on each test trial and present this as inputs to the model; we then evaluate the reconstruction error on those neurons alone. **Right:** In multiregion cosmoothing dropout training, we drop a percent of neurons from each region on some percent of training trials, and evaluate reconstruction error on all neurons.

### A.3 Double well simulation details

The simulation switches between two sets of dynamics, accumulation dynamics, and return dynamics. The system switches from accumulating to return at fixed time following the stimulus presentation and memory periods, mimicking the visual cue marking the end of the maze and the beginning of the maze choice arms for the animal in the real experiment.

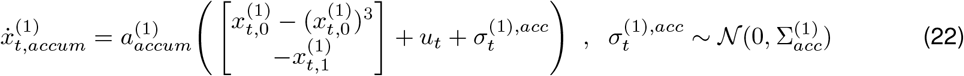

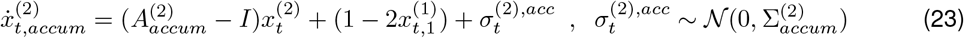

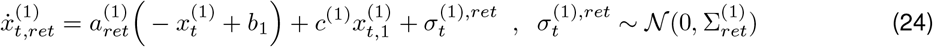

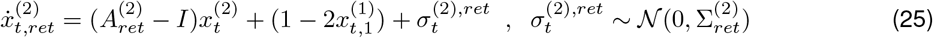

The following values are used for the variables:

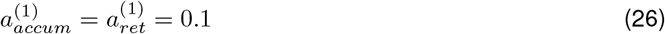

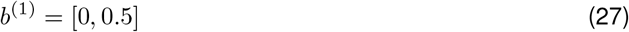

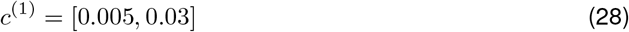

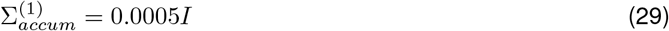

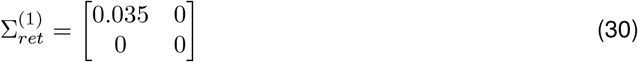

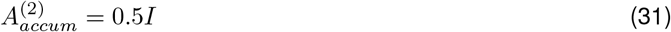

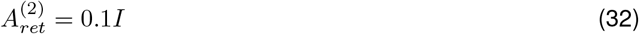

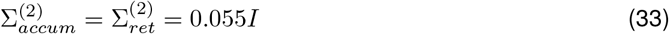

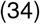

**Figure 7:**
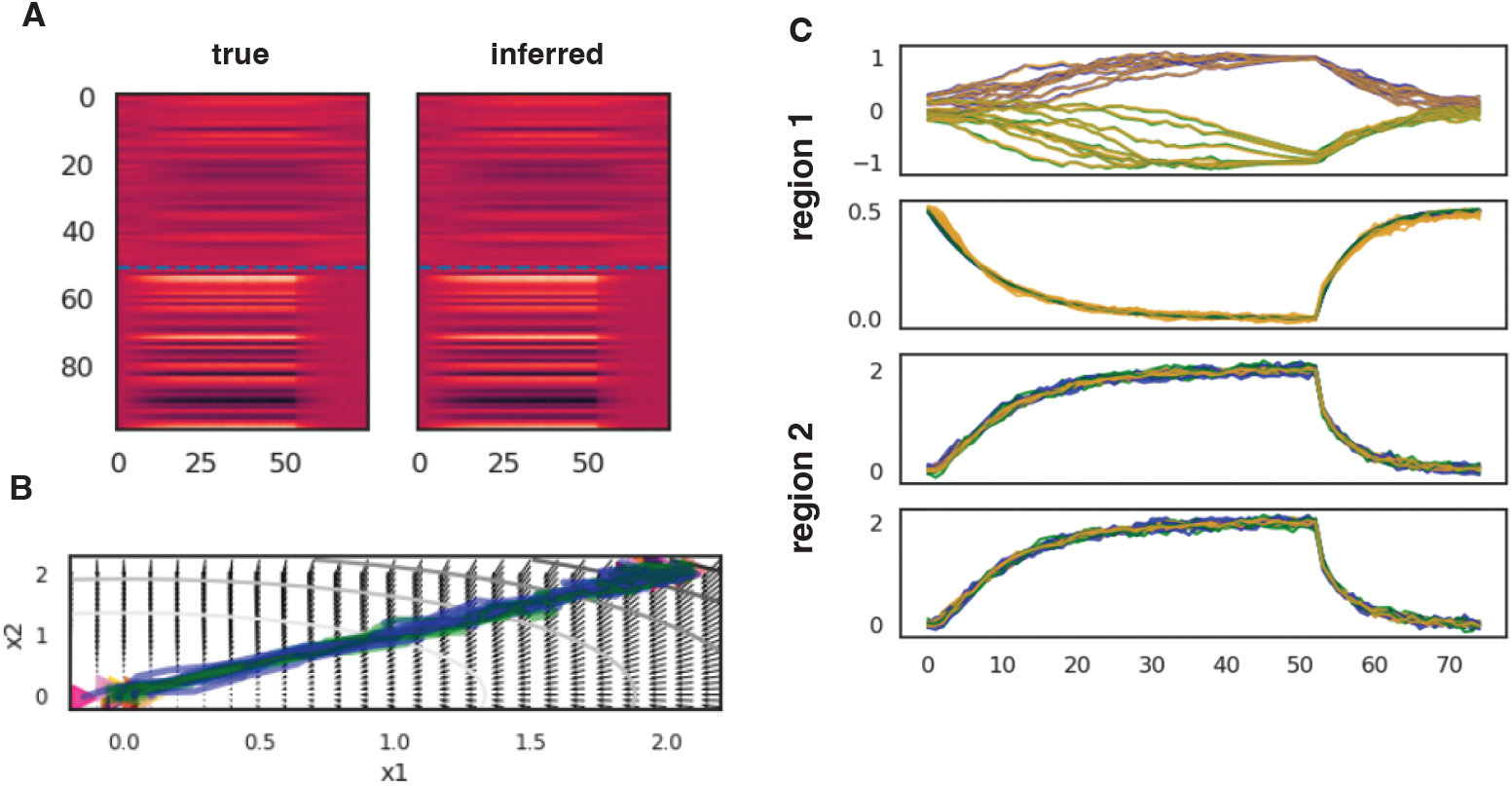
Double well simulation - additional details. **(A)** Recovery of observations for both regions. **(B)** Dynamics flow fields for controller (region 2) during accumulation period (qualitatively similar during return period). **(C)** Recovery of latents for both dimensions of both regions, across 18 test trials. Left and right trials in blue and green, recovered in yellow.

### A.4 Benchmarks

We provide details on the models used in the benchmarks presented across experiments.

#### A.4.1 MR-PCA

PCA is a surprisingly powerful baseline on many unsupervised machine learning tasks, e.g. (Bojanowski et al., 2017). We include in the benchmarks a comparison to multi-region PCA (MR-PCA), by which we mean fitting PCA to each region’s training data separately. Test data for each region is then reconstructed by first projecting it onto the top *d* principal components for that region. We note that while MR-PCA is a strong baseline for test, it can only compress data linearly, and this performs poorly on co-smoothing on both datasets, emphasizing the importance of this metric. Additionally, we note that MR-PCA is a much stronger baseline on the widefield dataset, which is consistent with the low intrinsic dimensionality of the data (see Figure A3 of cumulative variance explained for both datasets).

**Figure 8:**
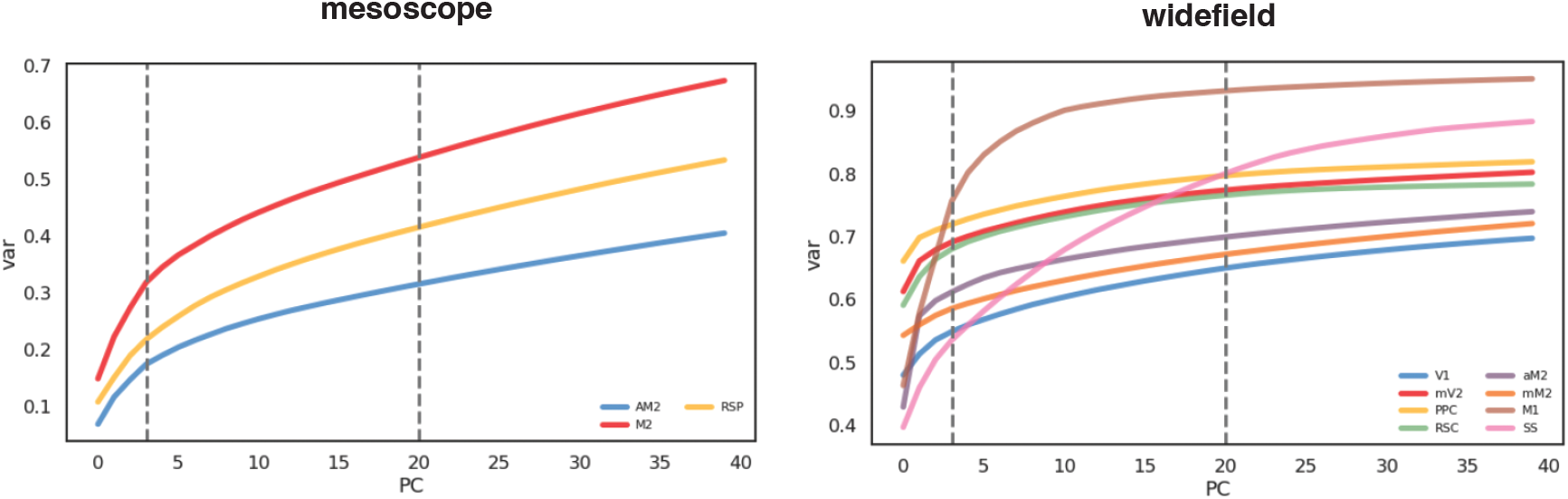
Variance explained by PC dimension per region, for both real datasets. Intrinsic (linear) dimensionality of the mesoscope data is higher than widefield.

#### A.4.2 Multiregion LDS, SLDS, and rSLDS models

Briefly, (Glaser et al., 2020) introduced a multiregion rSLDS model of the following form:

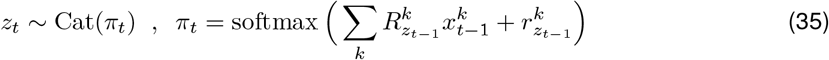

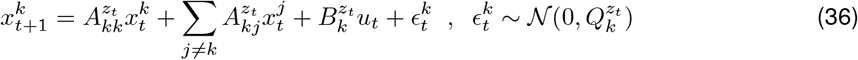

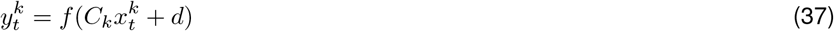

Where above *A, B, R, Q, C* are matrices and *r, d* are bias vectors. We refer to the model above as an MR-RSLDS. Similarly, we refer to versions of this model with 1 discrete state as an MR-LDS, and versions with no recurrence in the switching dynamics as MR-SLDS.

All LDS variant models were run in parallel on 28 CPUs using a modified version of the SSM package [Linderman et al https://github.com/slinderman/ssm], using variational Laplace EM (vLEM) (Zoltowski et al., 2020).

#### A.4.3 Higher-order auto-regressive, or AR(p) MR-RSLDS models

We include in the benchmarks comparison LDS and rSLDS models with higher order auto-regressive continuous latent dynamics. These models (AR(p) MR-RSLDS) have the following modified latent dynamics:

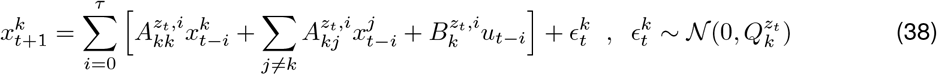

Above, *τ* is the order of the AR process, or the total number of lags, with *i* indexing the *τ* lagged dynamics, communication, and inputs matrices, 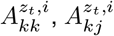 and 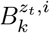. Similarly, *x*_*t−i*_ and *u*_*t−i*_ represent the *i*’th lagged state and input at time *t*. The emissions and discrete latent transition dynamics remain as in the single lag models.

Adding higher order autoregressive dynamics adds to the expressivity of this class of models. While dynamics remain piecewise linear, they are no longer linear with respect to the previous timepoint alone. We note that these models performed better than single lag models (see results in Tables 1,A1).

In order to fit these models, we extended vLEM (Zoltowski et al., 2020) to higher order lags. Briefly, this involved modifying the Hessian to account for higher order terms.

#### A.4.4 Single region models

In the benchmarks, we include a comparison with single region LDS and rSLDS models that have a latent dimension equal to the sum of the latent dimensions used across regions in the multiregion models. Fitting a single region model to multiregion data results in a meaningful loss of interpretability, because there are no per-region latents, and no estimates of communication. However, these models in general can achieve lower test and cosmooth errors, since they are free to use latent dimensions to explain any part of the data across regions. As such, they give a baseline for performance on these datasets. Table 1 shows that MR-SDS achieves lower cosmoothing test error than both single region LDS and rSLDS models on both datasets, thus providing an increase in both performance and interpretability. Table 2 shows that a better calibrated model improves in test error as well relative to single region LDS and rSLDS models on mesoscope data, but not widefield data. We think this is likely due to the higher intrinsic dimensionality of the mesoscope data (see earlier comments on dimensionality in the MR-PCA section). x

**Table 2:** Held-out test error with and without multiregion dropout training.

| dataset | $p_1$ | test | cosmooth |
| --- | --- | --- | --- |
| mesoscope | 0.8 | 0.0040009 | 0.0040761 |
| mesoscope | 0 | 0.0040569 | 0.0045115 |

### A.5 Experiment parameter details

We provide additional details on the network architecture and hyperparameters used to fit model for each experiment.:

**Table 3:** Parameters used for each of the experiments presented in the paper. LV: Switching Lotka-Voltera. DW: Double well. Meso: mesoscope. Wide: widefield. lr: learning rate. *p*_2_: trial dropout rate used for cosmoothing training. *I*: Identity mapping. L: linear layer. *d*: dimensionality of latents for each region. For Bi-RNN, semicolon (;) indicates a single layer with two concatenated RNNs of that size, one forward and one backward.

| | $K$ | $R$ | $d$ | $f_{kk,kj}$ | $f_{ku}$ | $g_k$ | $g_{emb}$ | $g_{birnn}$ | $g_{rnn}$ | $g_x^{\mu,\Sigma}$ | $p_1$ | lr | steps |
| --- | --- | --- | --- | --- | --- | --- | --- | --- | --- | --- | --- | --- | --- |
| LV | 2 | 2 | 1 | 32x2 | - | L | - | 3;3 | 6 | 64,32 | 0 | 1e-3 | 2.5e3 |
| DW | 2 | 2 | 3 | 32x2 | L | L | 16 | 3;3 | 6 | 32x2 | 0 | 1e-3 | 5e3 |
| Meso | 2 | 3 | 3 | 32x2 | 4x2 | 128x2 | 32x2 | 32;32 | 64 | 256 | 0.8 | 1e-3 | 15e3 |
| Wide | 2 | 8 | 2 | 32x2 | 4,2 | 32x3 | 32,16x2 | 32;32 | 64 | 128 | 0.5 | 2e-4 | 15e3 |

### A.6 Explicit duration

In the explicit duration case, the full generative model is:

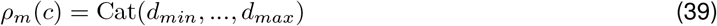

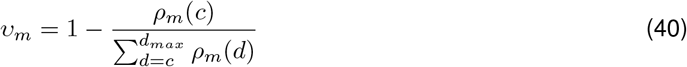

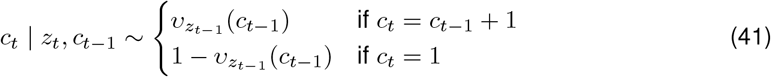

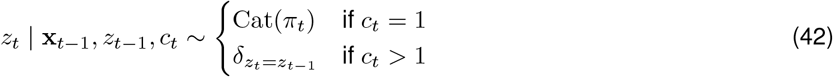

In words, each state *m* has a learned categorical distribution *ρ*_*m*_(*c*) over durations. Counts at time *t* are then sampled from a normalized version of this distribution, according to the discrete state *m* active at time *t* and denoted by *z*_*t*_. Conditioning on the counts allows the discrete state to persist without sampling a new state through the transition distribution *π*. The choice of hyper parameters *d*_*min*_, *d*_*max*_ confers an inductive bias on the augmented model, that impacts the persistence of inferred discrete states.

**Figure 9:**
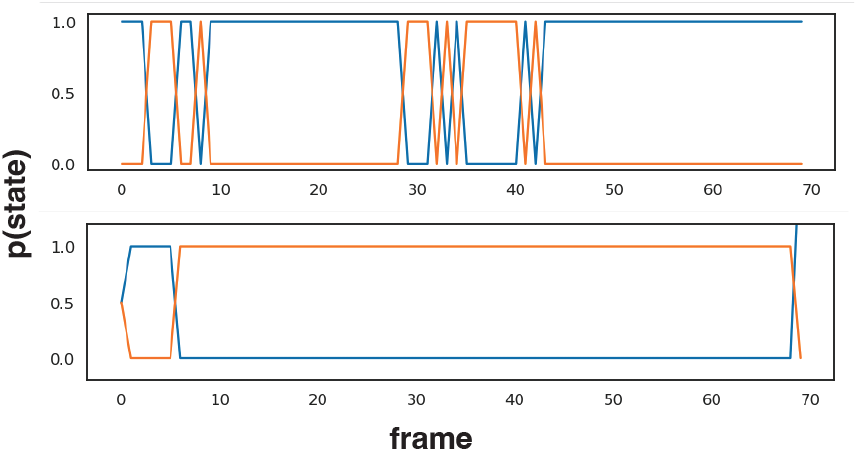
Explicit duration results in longer state persistence. **Above** Inferred discrete state for single mesoscope trial with no learned explicit duration variable. **Bottom:** Same, with explicit duration. Note the inferred discrete state persists for longer.

